# Adverse Effects of Culture Media on Human Pluripotent Stem Cells

**DOI:** 10.1101/067868

**Authors:** Megha Prakash Bangalore, Odity Mukherjee, Syama Adhikarla, Mitradas M. Panicker

**Affiliations:** National Centre for Biological Sciences (TIFR), Bangalore - 560065, India; Manipal University, Madhav Nagar, Manipal, Karnataka 576104; Institute for Stem Cell Biology and Regenerative Medicine, Bangalore 560065, India

**Keywords:** HPSCs, culture media, oxidative stress, nuclei acid damage, anti-oxidants, nucleoli

## Abstract

Culture conditions play an important role in regulating the genomic integrity of HPSCs. We report that HPSCs cultured in Essential 8 (E8) and mTeSR, two widely used media for off-feeder culturing of HPSCs, had many fold higher levels of ROS and higher mitochondrial potential than cells cultured in KSR containing media. HPSCs also exhibited increased levels of 8-hydroxyguanosine, phospho-histone-H2A.X and p53, as well as increased sensitivity to γ-irradiation in these two media. HPSCs in E8 and mTeSR had increased incidence of alterations in their DNA sequence, reflecting genotoxic stress, in addition to changes in nucleolar morphology and number. Supplementing E8 and mTeSR with antioxidants provided only a partial rescue. Our results suggest that it is essential to determine cellular ROS levels in designing culture media as it affects the genomic integrity of HPSCs and will limit their use in studying development and in regenerative medicine.

**Highlights:**

- Culture media can dramatically alter nuclear and nucleolar morphology
- HPSCs in E8 and mTeSR media have increased ROS levels and mitochondrial potential
- There is increased nuclei acid damage in HPSCs cultured in E8 and mTeSR media
- Nucleolar morphology of HPSCs can act as a “stress reporter”

## Introduction

The capability of human pluripotent stem cells (HPSCs) to self-renew as well as differentiate into all cell types make them valuable for therapy, in understanding early developmental processes and to model many human diseases. This unique property of stem cells is primarily attributed to and regulated by a number of complex and specialized processes. In order to fulfil their potential, it is essential that these cells maintain their unique properties as well as their genomic integrity. Various studies have indicated that the levels of reactive oxygen species in pluripotent stem cells (PSCs) are significantly lower than their differentiated counterparts (Armstrong et al., 2010; Cho et al., 2006; Saretzki et al., 2008). This has been hypothesized as a way to protect cellular components i.e. lipids, protein, RNA and DNA from oxidative damage. They are also reported to have increased abilities to repair their DNA to maintain genomic stability (Filion et al., 2009; Lin et al., 2014; Luo et al., 2012; Maynard et al., 2008; Momcilovic et al., 2010).

Over the years, several studies have aimed at making ‘clinically useful’ HPSCs. The source of somatic cells and the process of reprogramming have been examined to determine sources of genomic variation (Abyzov et al., 2012; Hussein et al., 2011; Ji et al., 2012; Raab et al., 2014). Extensive research has also gone into optimizing the ideal culture conditions to maintain and propagate HPSCs leading to the development of different substrates and media which are ‘chemically defined and xeno-free’, can support feeder-free cultures of HPSCs, show lower batch-batch variation and increased ease of handling (Amit, 2002; Chen et al., 2011; Hovatta et al., 2003; Li et al., 2005; Ludwig et al., 2006a, 2006b; Richards et al., 2002; Xu et al., 2001; Zhou et al., 2012). In the above studies, the ‘quality’ of stem cells has been defined by robust expression of pluripotency markers, capability to differentiate into all the three germ layers, established by teratoma formation or *in vitro* differentiation, and the presence of normal karyotypes after multiple passages. Efficient derivation of ESC and iPSC lines in these media has also been another criterion. Curiously, mitochondrial activity and reactive oxygen species (ROS) levels of established PSCs during routine culture due to the media *per se* have not been addressed. Perhaps, this has been, in part, due to early studies that have indicated that HPSCs depend on glycolysis and not on oxidative phosphorylation, and that PSCs, in general, exhibit low ROS levels (Armstrong et al., 2010; Mathieu et al., 2014; Saretzki et al., 2008; Zhang et al., 2012). A variety of media formulations are now available, many of them with antioxidants e.g. Vitamin C, but these again have not directly examined mitochondrial potential or ROS values of the cells cultured in them.

In an earlier study, we had identified lipid droplets containing retinyl esters as a marker unique to the ‘primed’ pluripotent state. We had also observed that these droplets were present in cells cultured in Knockout Serum Replacement (KSR) containing media but not in Essential 8 (E8) and mTeSR media (Muthusamy et al., 2014). This suggested that the metabolic state i.e. lipid metabolism, of HPSCs in these two media were different and led us to examine other aspects of HPSCs in these media, in more detail. We observed significant changes in the nuclear and nucleolar morphology of cells in the three media. Particularly, changes in the morphology of nucleoli which are known to be markedly affected by stress (Mayer and Grummt, 2005; Mayer et al., 2005; Westman and Hutten, 2010) led us to investigate the metabolic activity of HPSCs in different media which often impacts ROS and mitochondrial potential levels. Our study shows that HPSCs in E8 and mTeSR media have higher levels of ROS and mitochondrial potential when compared to KSR media. Associated with these, were higher levels of markers of double stranded DNA breaks (DSBs) and increased sensitivity to γ-irradiation induced DSBs. The RNA in HPSCs cultured in these two media also exhibited far more 8-hydroxy guanosine modifications, including increased levels in the nucleoli. The increased oxidative stress seen in E8 and mTeSR media would certainly affect their long term culture and genomic status. Associated with the higher ROS levels were also increased number of single nucleotide variations (SNVs) in the genomic DNA.

Most studies have assumed, the media commonly used to culture PSCs to be equivalent and differing primarily in their ease of use, number of components and their xeno-free status. Since the media used for culturing HPSCs can have very pronounced metabolic effects (Zhang et al., 2016), it would be necessary to also examine ROS levels and mitochondrial potential in addition to pluripotency markers and karyotypic changes in the process of designing culture medium for HPSCs. While karyotypic changes would report large changes in genomic DNA and have been used as a surrogate for genomic integrity, SNVs due to the media, have not been considered.

## Results

### Nuclei and nucleoli of HPSCs cultured in E8 and mTeSR media are morphologically distinct from those in KSR medium

HPSCs cultured in E8 and mTeSR media exhibited very different nuclear and nucleolar morphologies from those seen in KSR. This was also observed in images from the literature (Bogomazova et al., 2015) and from those provided by the manufacturers (http://www.stemcell.com/en/Products/Popular-Product-Lines/mTeSR-Family.aspx). The differences in the size and shape of cells in the three media were significant and also reproducible (Figures 1A, 1B and 1C). A systematic analysis using nuclear (Hoechst 3342) staining showed that while the nuclei in KSR media appeared larger, the nuclei in E8 and mTeSR were smaller. This was even more evident on examining the nuclear cross-sectional area which was the highest in KSR as compared to E8 and mTeSR (Figures 1A and 1C) even though the total nuclear volume in all three media remained constant (Figure 1C). Further analysis determined that the height of nuclei was the least in KSR and significantly higher in E8 and mTeSR (Figure 1C). HPSCs in KSR had a more elliptical/lobular nuclear cross-section, while in E8 and mTeSR, it was more circular as determined by the eccentricity values of the nuclear cross-sections (Figure 1C). Overall, HPSCs in E8 and mTeSR were smaller and more compactly arranged than in KSR, which was also evident from the number of nuclei per unit area of culture surface (Figure 1C).

**Figure 1:**
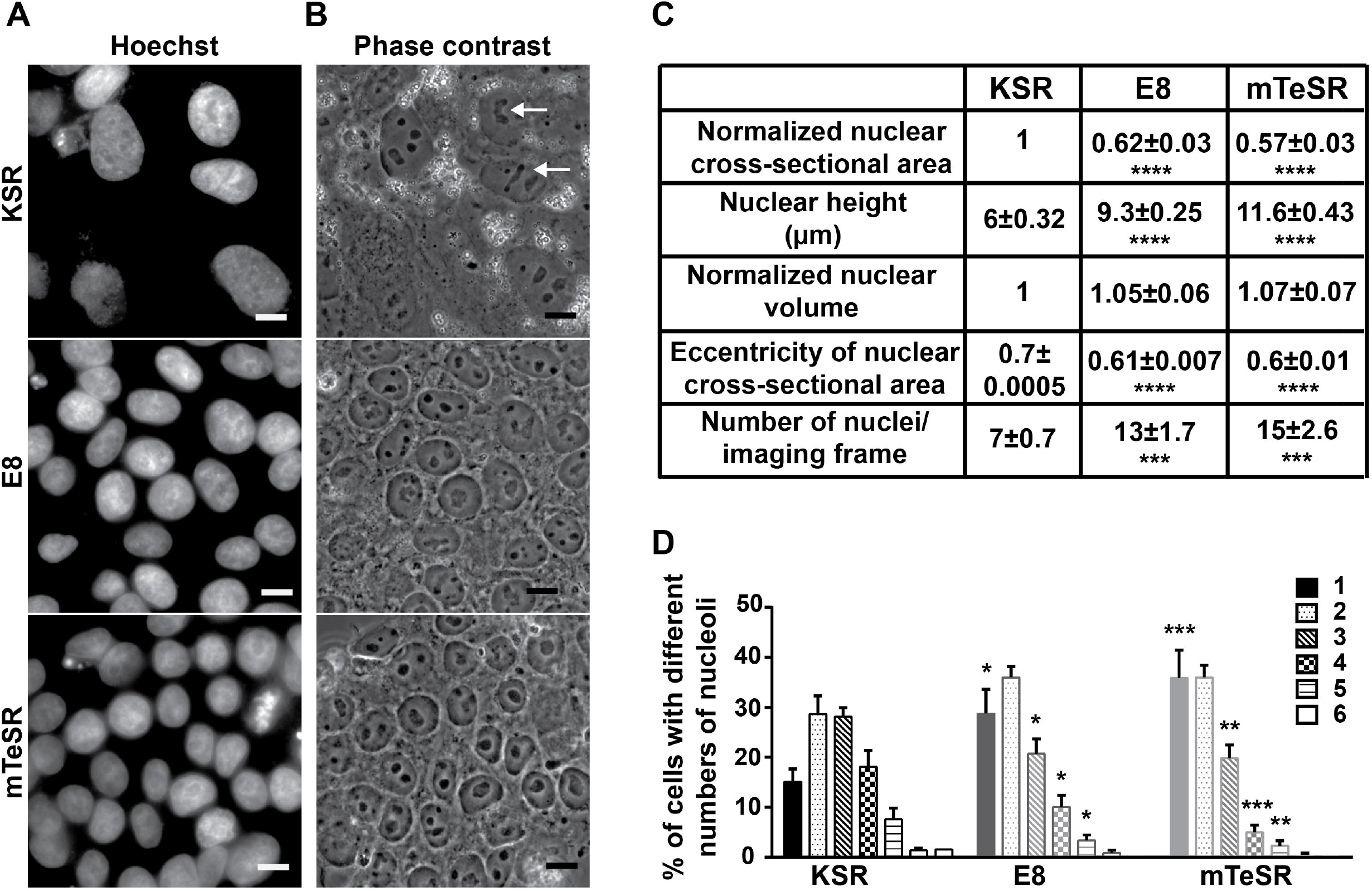
Nuclear and nucleolar morphologies of HPSCs cultured in KSR, E8 and mTeSR media are different. (A) Hoechst 33342 staining of ADFiPS showing larger nuclei in KSR compared to E8 and mTeSR media. (B) Bright field images of ADFiPS corresponding to their respective Hoechst images in (A) showing reticulate & multiple nucleoli per nucleus in KSR, fewer reticulate nucleoli and higher numbers of single nucleoli per nucleus in E8; large and round, mostly single nucleolus per nucleus in mTeSR. (C) Summary of various parameters defining the nuclei of HPSCs in the three media, (E) Quantification of the percentage of nuclei with different number of nucleoli for HPSCs in the three media. Normalized data represent values normalized with respect to KSR. Scale bars represent 10μ. Pooled data from all the four cell lines represented as mean ± SEM. Student’s t-test. **** p<0.0001, *** p<0.001, ** p<0.01, * p<0.05. n≥10 for all quantitative data.

HPSCs in these three media also exhibited differences in nucleolar morphology and numbers. In mTeSR, HPSCs predominantly had one or two large circular nucleolus/nucleoli per nucleus while in KSR, multiple reticulate nucleoli per nucleus were seen (Figures 1B and 1D). Cells in E8 also had increased numbers of nuclei with 1–2 predominantly circular nucleoli (Figures 1B and 1D). Reticulate nucleoli were the highest in KSR followed by E8 and almost completely absent in mTeSR (Figure 1B). These differences were most striking when cells at the periphery of a colony were observed as the distinction between individual cells were clearer; however, even at the centre of a colony where the cells were more tightly packed, these differences were apparent though less striking.

E8 and mTeSR have been in use for over five and ten years respectively, but surprisingly, these characteristics have not been examined earlier.

### HPSCs cultured in E8 and mTeSR have substantially higher ROS levels and mitochondrial potential

Nuclear morphology has been related to gene expression (Koh et al., 2011) and the nucleolus is known to be a stress sensor. Since there were pronounced media-dependant changes in the nucleolar morphology and numbers, we determined if the cells were stressed to different extents in the three media. We found that the ROS levels in HPSCs, which at high levels are a major inducer of stress, were dependent on the media. The ROS levels, measured through DCFDA fluorescence, were approximately 5 and 3 fold higher in E8 and mTeSR respectively, than in KSR (Figure 2A). High ROS values were seen within 48 hours of shifting cultures from KSR to E8 and mTeSR (Figure 2B). Interestingly, ROS levels of HPSCs cultured in E8 or mTeSR for three or more passages, when placed back in KSR did not decrease to the levels typically seen in KSR even after five to eight days (Figure 2C). Our data suggests that HPSCs in E8 and mTeSR are subject to higher ROS levels than in KSR and that the ROS levels are not easily reversed. This suggests that substantial changes in the underlying metabolic metabolism occurs in these two media which take time to revert.

**Figure 2:**
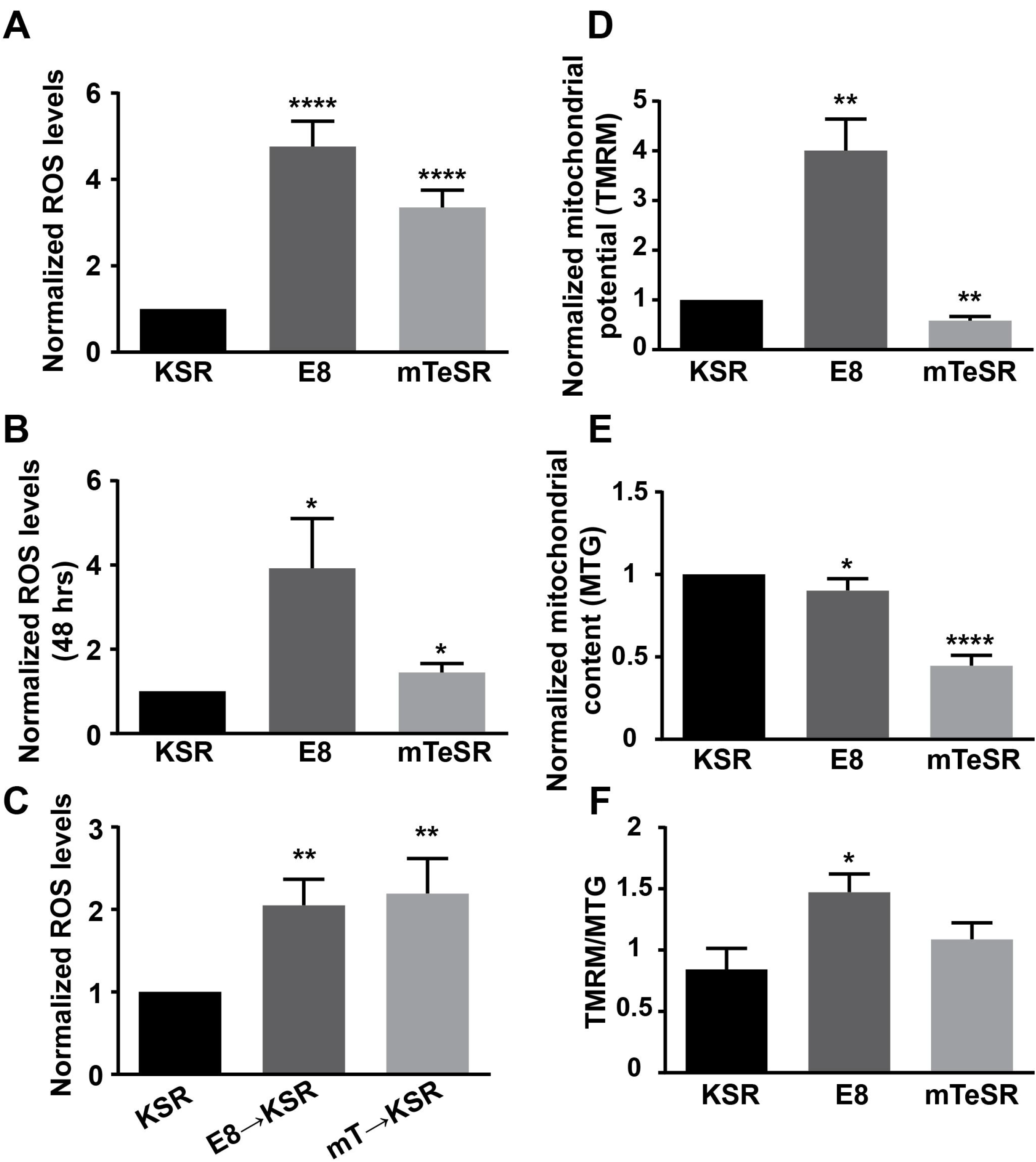
ROS and mitochondrial levels of HPSCs in E8 and mTeSR media are higher than in KSR. Quantification of DCFDA fluorescence values (ROS levels) of HPSCs cultured in the three media for (A) ≥3 passages (B) 48 hours and (C) in KSR media for 5–8 days after three passages in E8 and mTeSR; n≥10, n=3 and n=6, respectively. Quantification of mitochondrial potential expressed as (D) TMRM fluorescence/cell and (F) TMRM/MTG fluorescence; n≥5 and n=4 respectively, (E) mitochondrial content - MTG fluorescence; n=7. Normalized data represent values normalized with respect to KSR. Pooled data from all the four cell lines represented as mean ± SEM. Student’s t-test. **** p<0.0001, *** p<0.001, ** p<0.01, * p<0.05.

The high ROS levels led us to examine mitochondrial metabolism in these media. HPSCs are primarily dependent on glycolysis. Even with well-developed mitochondria, they do not provide ATP through oxidative phosphorylation due to high levels of UCP2, an uncoupling protein (Zhang et al., 2011), and have lower mitochondrial potentials. We examined the mitochondrial potential of HPSCs in the three media. The mitochondrial potential (TMRM fluorescence) of HPSCs was ~3.7 fold higher in E8 and ~0.5 fold lower in mTeSR when compared to that in KSR (Figure 2D).

We also examined the mitochondrial content (MTG fluorescence) of HPSCs in the three media (Figure 2E). HPSCs in mTeSR had ~half the mitochondrial content when compared to KSR and E8, which in turn showed marginal differences. However, if the mitochondrial potential was normalized to mitochondrial content, it was higher in mTeSR as compared to KSR although not statistically significant (Figure 2F). However, mitochondrial potential computed as TMRM fluorescence per cell and TMRM fluorescence per unit mitochondrial mass, were both higher in E8 compared to KSR (Figures 2D and 2F).

Our data suggests that the mitochondria of HPSCs are metabolically more active when cultured in E8 and mTeSR than in KSR, as measured by mitochondrial potential.

### Altered ROS levels in HPSCs in E8 and mTeSR results in increased nucleic acid damage

ROS generated as a by-product of cellular metabolism is a major source of DNA damage. Since HPSCs in E8 and mTeSR had higher ROS levels, we measured the extent of oxidative damage in these cells by probing for nuclear 8-hydroxyguanosine (8OHG). HPSCs in mTeSR had the highest levels of nuclear 8OHG followed by E8 while cells in KSR had negligible levels (Figures 3A and S1A). Thus, higher levels of ROS in E8 and mTeSR were indeed associated with increased oxidative damage. Double stranded breaks (DSBs), which are known to occur at a very low frequency in routine cultures, were also substantially higher in cells cultured in E8 and mTeSR (Figure 3B) measured by γ-H2AX immunofluorescence. Additionally, sensitivity to γ-irradiation, was also higher in HPSCs cultured in E8 and mTeSR when compared to KSR (Figure 3C). Furthermore, p53 which is known to accumulate in the nucleus in response to DNA damage in HPSCs (Gambino et al., 2013; Grandela et al., 2008) was observed to be higher in the nuclei of HPSCs cultured in E8 and mTeSR when compared to that in KSR (Figure 3D).

**Figure 3:**
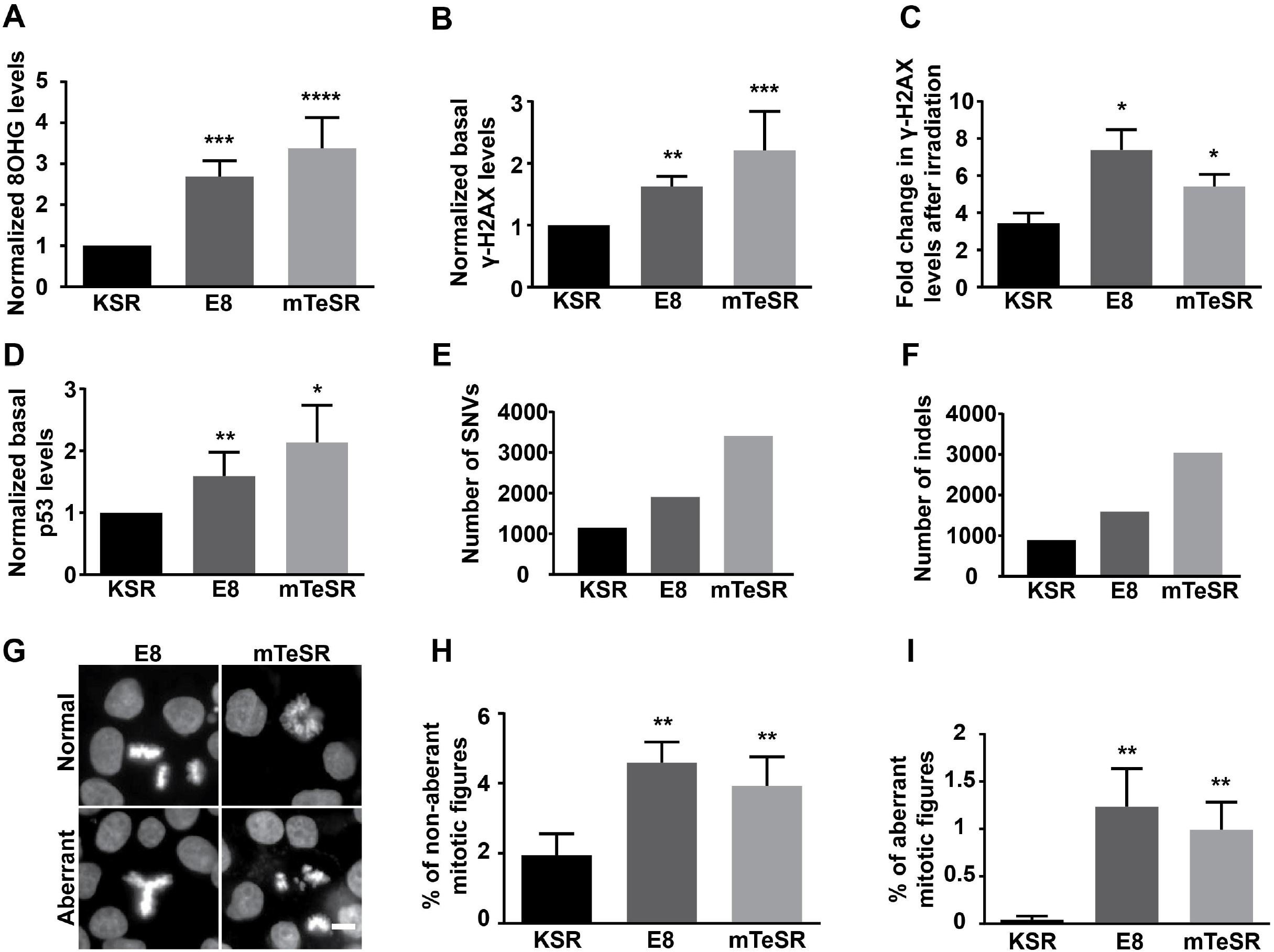
HPSCs in E8 and mTeSR media show higher nuclei acid damage when compared to KSR. Quantification of (A) normalized nuclear 8-hydroxyguanosine levels; n=9, (B) basal γ-H2AX levels – DSBs; n≥5, (C) fold increase in DSBs after γ-irradiation; n≥3, (D) normalized basal p53 levels; n≥4, of HPSCs in the three media. (E) Total number of SNVs, (F) insertions and deletions seen in HuES9 cells cultured in the three media for three passages. (G) Representative images and quantification of percentage of (H) non-aberrant mitotic figures and (I) aberrant mitotic figures of HPSCs in the three media; n=12. Normalized data represent values normalized with respect to KSR. Scale bars represent 10μ. Pooled data from all the four cell lines represented as mean ± SEM. Student’s t-test. **** p<0.0001, *** p<0.001, ** p<0.01, * p<0.05.

Exome sequencing data from HuES9 cells cultured in the three media showed increased number of SNVs in E8 (1.7 times) and mTeSR (3 times) when compared to KSR (Figure 3E). The total number of insertions and deletions were also higher in E8 (1.4 times) and mTeSR (3.4 times) than in KSR (Figure 3F).

Higher numbers of mitotic figures were seen in HPSCs in E8 and mTeSR than that in KSR (Figure 3G and 3H). This may reflect the higher proliferation rate of HPSCs in E8 and mTeSR reported in the literature (Chen et al., 2011) or it may be due to more cells undergoing mitotic arrest in E8 and mTeSR due to DNA damage. In addition to increased number of mitotic figures (non-aberrant), the number of aberrant mitotic figures were also higher in E8 and mTeSR (Figure 3G and 3I). Representative aberrant mitotic figures observed in E8 and mTeSR are shown in Figure S1B and S1C.

These observations suggest that HPSCs in E8 and mTeSR are not only subjected to increased basal DNA damage but are also more sensitive to genotoxic treatments e.g. γ-irradiation.

### Addition of antioxidants to E8 and mTeSR provides partial rescue from genotoxic stress

Higher levels of ROS and increased mitochondrial potential in E8 and mTeSR associated with increased DNA damage led us to investigate the effect of increasing the levels of antioxidants in the media. Glutathione which is absent in E8, and Vitamin C - two commonly used antioxidants in HPSC culture were used (Ji et al., 2014; Luo et al., 2014). ROS levels decreased significantly in E8 and mTeSR with the addition of antioxidants but not to the levels observed in KSR (Figure 4A). Mitochondrial potential however, was not affected with the addition of antioxidants (Figure 4B).

**Figure 4:**
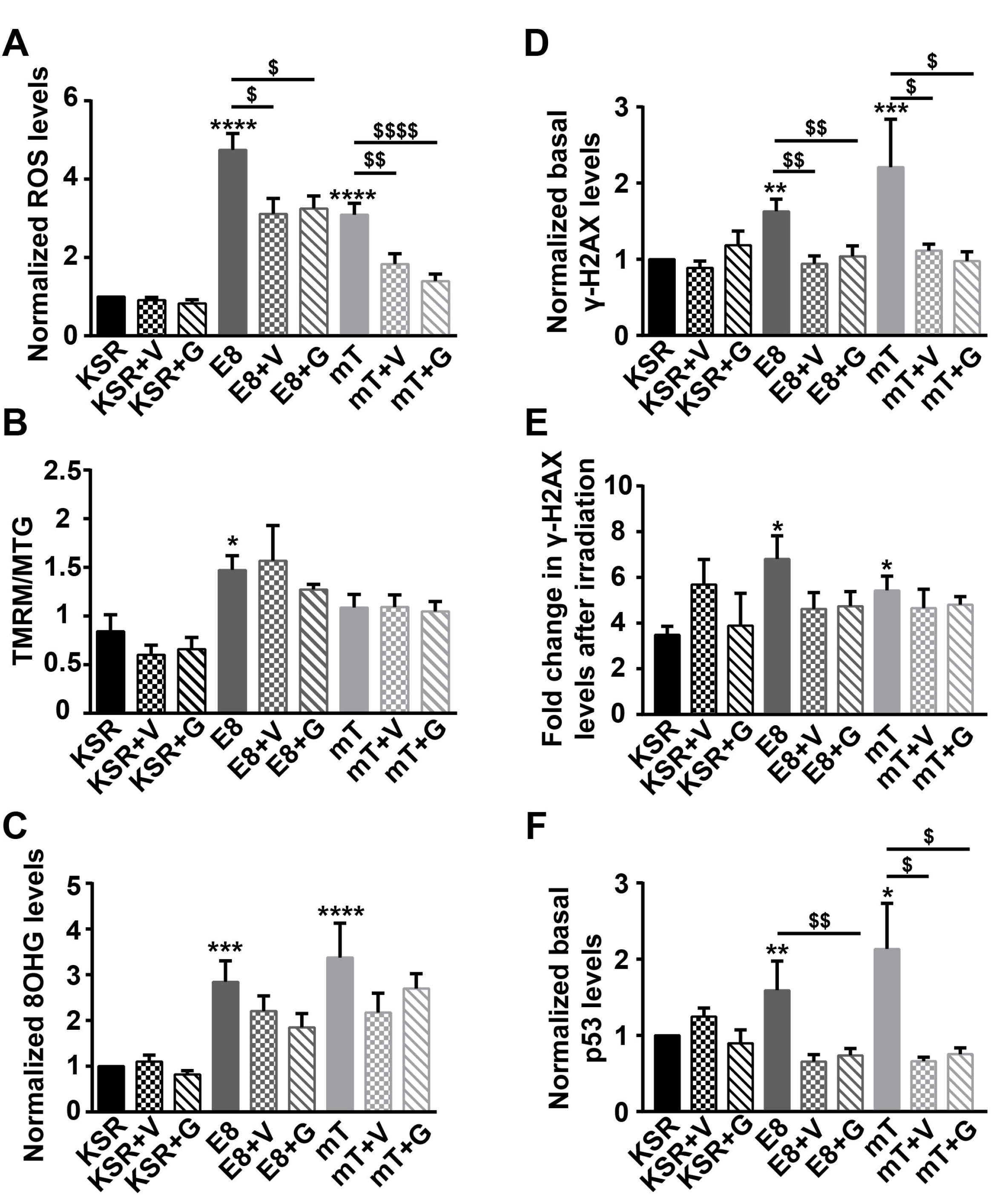
Vitamin-C and glutathione (GSH) partially reduce the effects of E8 and mTeSR media on HPSCs. Quantification of (A) normalized ROS levels; n≥10, (B) normalized mitochondrial potential levels; n=4, (C) normalized nuclear 8OHG levels; n=6, (D) normalized basal DSBs (γ-H2AX); n≥5 (E) fold increase in DSBs after γ-irradiation; n=4 and (F) normalized p53 levels; n≥4 for HPSCs cultured in the three media. Normalized data represent values normalized with respect to KSR. V: Vitamin-C, G: Glutathione. Pooled data from all the four cell lines represented as mean ± SEM. Student’s t-test. **** p<0.0001, *** p<0.001, ** p<0.01, * p<0.05. $ represent p-values with respect to their untreated controls.

8OHG levels also decreased with the addition of antioxidants but the values were not statistically significant and never reached the levels seen in KSR (Figure 4C). Basal γ-H2AX levels however, showed significant decrease with antioxidants and reached values similar to that of KSR (Figure 4D). Sensitivity to γ-irradiation showed a decreasing trend with the addition of antioxidants to E8 and KSR but the values were not statistically significant (Figure 4E). Basal p53 values, showed both substantial and significant decrease (Figure 4F). These results suggest that higher levels of ROS in E8 and mTeSR were responsible for increased DNA damage in HPSCs and that addition of antioxidants was only partially effective.

### Nucleolar morphology can act as an indicator of genotoxic stress

HPSCs in E8 and mTeSR, with higher ROS levels and associated nuclei acid damage than cells in KSR, also possess distinct nucleolar morphologies i.e. rounded and non-reticulate nucleoli unlike in KSR. Since nucleoli are known to be ‘stress sensors’, we examined the change in nucleolar morphology of HPSCs cultured in KSR after subjecting them to genotoxic stress. On treatment of HPSCs in KSR with doxorubicin for three hours, 70.4% of nuclei showed distinct and rounded nucleoli (Figures S2A and S2B) as opposed to only 22.6% in untreated HPSCs (Figure S2C). HPSCs exposed to γ-irradiation also showed a higher proportion of nuclei with distinct and rounded nucleoli (42.2%) within an hour (Figure S3). Thus, nucleolar morphology in HPSCs does respond to genotoxic stress and within short time frames. However, the number of nucleoli per cell did not decrease dramatically with these treatments unlike its shape.

At the concentrations of the antioxidants used, we also did not observe a reversal of the nucleolar morphology of HPSCs in E8 and mTeSR to that in KSR, reinforcing the observation that even with increased levels of antioxidants in E8 and mTeSR, the rescue from oxidative stress was only partial. This may be due to ROS values of HPSCs cultured in E8 and mTeSR remaining high even after being transferred back to KSR media for five-eight days.

## Discussion

The culture of HPSCs for regenerative purposes and as model systems to study development and disease, rests on the genomic integrity of the cells through repeated passages. Our initial observations that HPSCs cultured in E8 and mTeSR exhibit differential lipid metabolism and are strikingly different, morphologically, from those cultured in KSR led us to compare HPSCs grown in these three media. The choice of media that we have examined in this study was, in part, dictated by their widespread use. HPSCs grown in E8 and mTeSR have become increasingly popular because of their limited, defined and xeno-free components. These media have been tested through multiple passages using a few criteria i.e. stable expression of pluripotency markers, ability to differentiate into all three germ layers (either *in vitro* or through teratoma formation), stable karyotype and also permissiveness in deriving embryonic and induced pluripotent stem cells. Pluripotent stem cells (PSCs) are reported to have low levels of reactive oxygen species (ROS), and the reduced environment seen in these cells has been attributed as a protective mechanism to prevent nucleic acid, protein and lipid damage but the impact of the various media on ROS levels within these cells have not been compared.

Our results indicate that the ROS levels of HPSCs in E8 and mTeSR are many fold higher than in KSR. The mitochondrial potential is also higher in HPSCs when grown in these two media. High levels of ROS are understandably detrimental to the integrity of cells, except in cases where small increases in ROS are known to be a signal for replication (Schmelter et al., 2006). The effects of high levels of ROS and associated DNA damage were evident in cells cultured in E8 and mTeSR with increased number of changes in their DNA sequence, presence of a small but significantly higher percentage of cells with mitotic figures and cells being more prone to genomic damage when subjected to radiation. The increased numbers of mitotic figures in HPSCs cultured in E8 and mTeSR may represent ‘arrested’ nuclei in response to high levels of ROS and subsequent DNA damage and remains to be investigated further. A recent study which examined anueploid HPSCs showed that due to segregation errors, higher percentage of cells exhibited ‘trailing’ DNA during mitosis as opposed to diploid HPSCs (Golan-lev et al., 2016). Interestingly, such abnormalities were also evident in our HPSCs cultured in E8 and mTeSR media but not in KSR (Figures 3G, 3I, S1B and S1C). All these observations strongly suggest that cells in E8 and mTeSR media are under severe genotoxic stress and perhaps have defective replication dynamics.

Since PSCs give rise to differentiated somatic cells as well as form germ cells, they are expected to be protected from genotoxicity. However, they should also undergo cell death, when the damage is substantial. Both mouse and human PSCs have been reported to have more robust oxidative stress defence mechanisms than their differentiated counterparts. So, it is likely that they can survive genotoxic stress up to certain levels beyond which their genomic integrity would be too compromised. This is perhaps what we observe in E8 and mTeSR where HPSCs are subjected to high ROS levels but not high enough that they cease to replicate. Though resistant to ROS levels, HPSCs show increased sensitivity to DNA damage, perhaps due to the DNA repair being over-extended even at a basal level. This is reflected by the increase in single nucleotide variations and aberrant mitotic figures seen in HPSCs in E8 and mTeSR. Moreover, HPSCs grown in E8 or mTeSR, show increased levels of DNA damage markers, namely, γ-H2AX, nuclear p53 and 8-hydroxyguanosine compared to KSR. Cells cultured in E8 and mTeSR for multiple passages are reported to have normal karyotypes (Chen et al., 2011; Ludwig et al., 2006b) but that does not exclude the presence of smaller changes in the genome.

A striking but unreported feature that we noticed in HPSCs cultured in these three media were their differences in nuclear and nucleolar morphology. In E8 and mTeSR, the nuclei of HPSCs are more compact and appear smaller in cross-section while HPSCs in KSR mainly have lobular nuclei and also occupy a significantly larger area. Another striking feature was the difference in the number and shape of the nucleoli of HPSCs cells in these three media as described earlier (Figures 1B and 1D). Nucleoli are known to be stress sensors, since they regulate translation and are known to get altered due to stress. On stressing HPSCs grown in KSR, with γ-irradiation or doxorubicin i.e. genotoxic stress, the reticulate and dispersed nucleoli become circular and more distinct. This is in accordance with HPSCs being under greater genotoxic stress in E8 and mTeSR. The observation that the number and shape of the nucleoli can report on cellular stress levels is useful and can be used to monitor the health of HPSCs.

Interestingly, the ROS levels remained higher when cells from E8 and mTeSR were transferred back to KSR even after five-eight days. It is likely that, with longer periods in KSR, the ROS levels may decrease to the levels seen in cells in continuous culture in KSR. However our results indicate that the effect of media on the process of ROS generation and metabolism, in general, can be long-lasting. An increase in the number of genomic changes - SNVs and aberrant mitotic figures, are of concern when propagating cells in similar media, even for short durations or for derivation of new lines (Ji et al., 2014). While these media possess the advantage of having defined components and are feeder-free and xeno-free, our results indicate that they can compromise genomic integrity during cell expansion.

Antioxidants have been used as supplements in various media for culturing HPSCs. While E8 contains Vitamin C, mTeSR contains both glutathione and Vitamin C. In spite of this, HPSCs in these two media had high ROS levels and increasing the concentration of antioxidants had only a partial effect on decreasing ROS levels. It is not apparent at the moment, why the ROS levels are high in these two media and modifications will require more careful examination. It is clear from the lack of lipid droplets in HPSCs cultured in E8 and mTeSR that lipid metabolism of HPSCs is also different in the three media. HPSCs cultured in KSR and E8 have also been shown to have dramatically different lipid metabolism, very recently (Zhang et al., 2016). There have been quite a few studies which have compared different media for culturing HPSCs (Benvenisty et al., 2010; Chin et al., 2010; Denning et al., 2006; George et al., 2009; Hovatta et al., 2003; Marinho and Muotri, 2015; Nakagawa et al., 2014; Ramos-Mejia et al., 2012) but almost every study focused on the levels of pluripotency markers, karyotypic changes and the differentiation capabilities of HPSCs in the different media to evaluate their relative merits. A very recent study on the effect of cell numbers on genomic stability (Jacobs et al., 2016) established that minor changes in culture conditions can affect the health and quality of HPSCs strongly. Our study explores the levels of reactive oxygen species, in particular, and shows that some very commonly used culture media have dramatic and deleterious effects on the health of HPSCs, and strongly suggests that ROS levels are important in designing new media.

HPSCs destined to be used for therapeutic treatments will require expansion and have to be free of genomic aberrations. Thus, it becomes crucial that the cells be tested for not only gross genomic abnormalities but in addition, for DNA damages such as SNVs and DSBs, which are not routinely tested for. These parameters need to be considered in the design of media for culturing HPSCs, especially for clinical use.

### Experimental procedure

|  |  |
| --- | --- |
| KSR media | 20% Knockout Serum Replacement (KSR) in Knockout DMEM supplemented with 5ng/ml bFGF (Peprotech), Glutamax, Non-Essential Amino Acids, 2-mercaptoethanol, Penstrep (All from Invitrogen) |
| E8 media | According to manufacturer's instructions (A1517001) |
| mTeSR media | According to manufacturer's instructions (05850) |

Cell culture: Human embryonic stem cells (HuES-7; HuES-9) and human induced pluripotent stem cells (adult dermal fibroblast iPS; neonatal foreskin fibroblast iPS) were cultured without feeders on Matrigel (Corning, #354277) for 1–3 passages before starting the experiments. 1:1 mixture of conditioned and unconditioned KSR media was used as ‘KSR’ on Matrigel. Cells were cultured on Vitronectin (Life Technologies, A14700) for E8 and Matrigel was used for mTeSR. Vitamin C (25 μg/ml) and glutathione (10μg/ml) were supplemented every day during media change. 0.5mM EDTA (Life Technologies, 15575020) was used to dissociate cells for regular passaging.

Measurement of ROS levels: 2’,7’-Dichlorofluorescin diacetate (Sigma, D6883) was used at 10μM. Verapamil hydrochloride (Sigma, V4629) was used at 5μM with DCFDA to prevent efflux of DCFDA as shown previously (Armstrong et al., 2010). Briefly, cells were incubated with the dye for 20 minutes at 37°C and 5% CO_2_ after which, they were harvested as cell suspension in 1X PBS for analyses using FACS VERSE. Propidium iodide (1μg/ml) was used for selecting live cell population.

Measurement of mitochondrial potential: Tetramethylrhodamine Methyl Ester Perchlorate - TMRM (Sigma, T668) was used at 10nM along with Mitotracker green (Molecular Probes M7514) at 150nM. Briefly, cells were incubated with TMRM for 15 minutes at 37°C and 5% CO_2_ following which, they were harvested as cell suspension in 1X PBS and analysed using BD FACS Aria. Images were acquired on Nikon Eclipse TE2000-E-PFS (Japan) microscope with a Photometrix Cascade II 512 EM-CCD camera (Roper Scientific, USA) and Image Pro-Plus AMS software (Media Cybernetics, USA).

Geo-mean values of the respective area histograms were used for all the FACS data analyses.

Immunostaining and microscopy: Antibodies used were Anti-8 Hydroxyguanosine (Abcam, ab62623; 1:300), Phospho-Histone H2A.X (Ser139) (Cell Signaling Technology, #2577; 1:300), p53 (sc-126; Santa Cruz Biotechnology, INC, 1:100). Alexa Fluor dyes were used as secondary antibodies. Images were acquired on Olympus FV 1000 confocal microscope and analysed using ImageJ and Cell profiler.

Exome sequencing and analyses: Genomic DNA was extracted from cells cultured in the three media after three passages by salting out method (Miller et al., 1988). DNA libraries were prepared as per manufacturer’s protocol (Nextra rapid capture kit). Pair end sequencing was performed on Illumina Hiseq 2000 platform. The fastq files generated were aligned with human reference genome, hg19 built37 using bwa ver 0.5.9-r16. Indel realignment was performed using GATK (https://www.broadinstitute.org/gatk/). SNP variants were called by Varscan ver 2.3.6 (Koboldt et al., 2009) with a coverage of at least 8 and p value of 0.001. These variants were annotated using variant effect predictor (McLaren et al., 2016).

## Author contributions

MMP and MPB conceived the study and wrote the manuscript. MPB performed the experiments and analyses. OM and SA did the exome sequencing and analyses.

## Acknowledgements

We thank Council for Scientific and Industrial Research (CSIR) for salary (MPB), Shanta Wadhwani Centre for Cardiac and Neural Research (SWCCNR) for funding (OM) and NCBS (TIFR) for funding, Dr. Annapoorni Rangarajan (IISc, Bangalore) for providing doxorubicin, Deepak Kumar (AIIMS, New Delhi) for help with DNA isolation for exome sequencing, Thangaselvam Muthusamy (NCBS) for help with TMRM flow cytometry experiments, Doris Santiago (NCBS) for help with image analyses, Maki Murata-Hori (In-Stem) for help with reagents and CIFF, NCBS where all the confocal microscopy and flow cytometry experiments were undertaken. We are grateful to all the MMP laboratory members for their valuable suggestions and discussions.

